# Irisin Directly Stimulates Osteoclastogenesis and Bone Resorption *In Vitro* and *In Vivo*

**DOI:** 10.1101/2020.05.09.085977

**Authors:** Eben G. Estell, Phuong T. Le, Yosta Vegting, Hyeonwoo Kim, Roland Baron, Bruce Spiegelman, Clifford J. Rosen

**Author notes:** Corresponding Author: Clifford J. Rosen. This manuscript is submitted to eLife as a **Short Report**.

## Abstract

The myokine irisin facilitates muscle-bone crosstalk and skeletal remodeling in part by its action on osteoblasts and osteocytes. In the current study we investigated whether irisin also directly regulates osteoclasts. *In vitro*, irisin (2-10 ng/mL) increased osteoclast differentiation in C57BL/6J bone marrow progenitors; this increase was blocked by a neutralizing antibody to integrin α_V_β_5_. Irisin also increased resorption on several substrates *in situ*. RNAseq revealed differential gene expression induced by irisin including upregulation of markers for osteoclast differentiation and resorption, as well as osteoblast-stimulating ‘clastokines’. *In vivo*, forced expression of the irisin precursor *Fndc5* in murine muscle resulted in low bone mass and increased number of osteoclasts. Taken together, our work demonstrates that irisin acts directly on cultured osteoclast progenitors to increase differentiation and promote bone resorption. These actions support the tenet that irisin not only stimulates bone remodeling but may also be an important counter-regulatory hormone during exercise.

## Introduction

Irisin is a peptide generated by proteolytic cleavage of fibronectin type III domain-containing protein 5 (FNDC5), a membrane-bound protein highly expressed in skeletal muscle. *Fndc5* expression increases in response to acute bouts of exercise under regulation by PGC-1α, leading to a burst of circulating irisin^1^. Initially irisin was described as a circulating hormone that induces thermogenesis in adipose tissue^2^, but more recent work has shown a potent ability to modulate bone turnover. These effects support the tenet that irisin may be a key mediator of muscle-bone crosstalk during exercise. Initial studies demonstrated that irisin enhanced cortical bone formation and prevented unloading-induced bone loss i*n vivo*, and stimulated osteoblasts *in vitro*^3–5^. Conversely, genetic deletion of *Fndc5* was separately shown to block resorption-driven bone loss and maintain osteocyte function following ovariectomy; irisin treatment *in vitro* also prevented osteocyte apoptosis and stimulated sclerostin and RANKL release, key promoters of osteoclastogenesis, through the α_V_β_5_ integrin receptor^6^. The present study addresses the hypothesis that irisin also directly stimulates osteoclast differentiation and function *in vitro* and *in vivo*.

## Results and Discussion

First, we used continuous treatment with increasing doses of irisin (0, 2, 5,10 ng/mL) for 7-days in an *in vitro* osteoclast differentiation assay using primary marrow hematopoietic progenitors. These doses were based on previous work establishing the physiologic range of circulating irisin during and after exercise^1^. We found a qualitative enhancement of both osteoclast number and size (Figure 1a), and significant increases in osteoclast number across this dose range (2 ng/mL, 5 ng/mL, 10 ng/mL: *P* < .0001, 20 ng/mL: *P* = .044) (Figure 1b). Based on these results, we selected 10 ng/mL irisin for further experiments using both continuous (7 day) and transient treatment for the first 4 or 24 hr of culture. Both transient treatments led to enhanced osteoclast numbers versus controls (4 hr: *P* = .0218, 24 hr: *P* = .0008) (Figure 1c), but continuous irisin resulted in the largest increase (*P* < .0001) and was higher than both 4 hr (*P* = .0002) and 24 hr only treatments (*P* = .0152). The stimulatory effect of continuous 10 ng/mL irisin was then further confirmed in primary hematopoietic cells from both sexes of C57BL/6J mice (*P* = .0003), and in the RAW 264.7 macrophage cell line (*P* = .0428) (Figure 1d). RAW-derived osteoclasts appeared morphologically similar to primary cells and mirrored observations of qualitatively larger cells with irisin treatment (Figure 1e).

**Figure 1.**
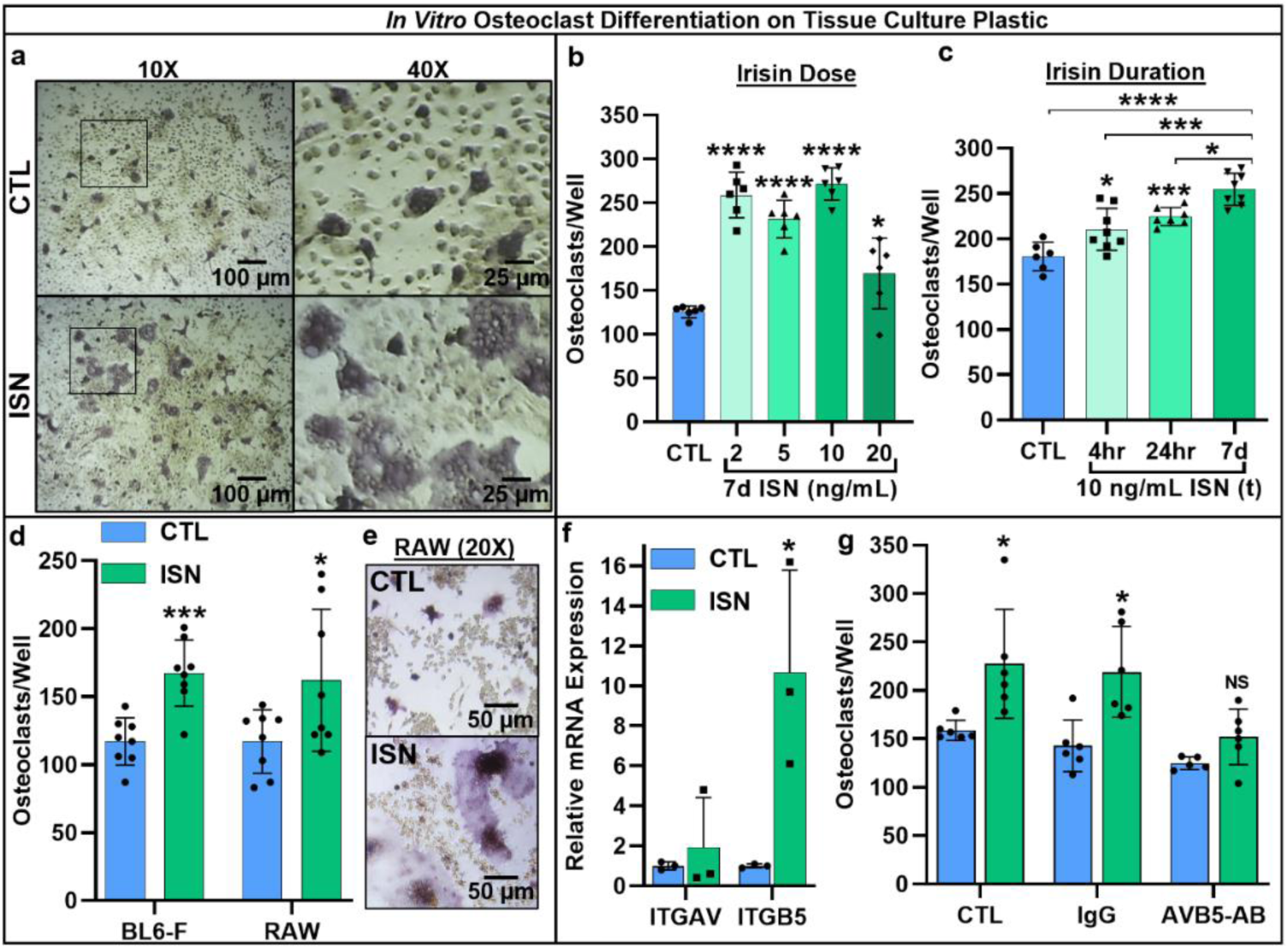
Representative 10X and 40X (boxed inset) images of TRAP-positive stained osteoclasts after 7-day differentiation with 10 ng/mL irisin (ISN) or untreated controls (CTL) (a). Quantification of osteoclasts per well demonstrating enhanced osteoclastogenesis in response to continuous (7d) ISN across a physiologic range (2-20 ng/mL) (b), and to treatment with 10 ng/mL ISN for only first 4 or 24 hr of culture compared to continuous treatment or CTL (c). Quantification of osteoclasts per well confirming irisin stimulation of osteoclastogenesis with continuous 10 ng/mL treatment across primary murine gender with female BL6 mice, and with the macrophage cell line RAW 264.7 (d), with representative images of differentiated RAW-derived osteoclasts (e). Expression of integrin receptor subunit αV (ITGAV) and β5 (ITGB5) in primary osteoclast cultures normalized to *Hprt* (f), and quantification of osteoclast per well counts for CTL or continuous 10 ng/mL ISN in the presence of integrin α_V_β_5_ neutralizing antibody (AVB5-AB), an IgG antibody control (IgG), or no antibody (CTL) (g). N = 3-8 wells/group, \**P* < .05, \*\**P* < .01, \*\*\**P* < .001, \*\*\*\**P* < .0001 vs. CTL within group or as indicated.

As integrins are found on the osteoclast membrane and known to play a role in differentiation^7,8^, and earlier work identified integrin α_V_β_5_ as a receptor for irisin on osteocytes^6^, we examined the expression of both subunits in osteoclast cultures and found increased relative mRNA expression above controls with irisin treatment (α_V_: *P* = .552, *β*_5_: *P* = .031) (Figure 1f). Blocking integrin α_V_β_5_ with a neutralizing antibody (AVB5-AB) resulted in no differences in osteoclast number per well for irisin treatment (10 ng/ml) versus untreated controls (*P* = .79) compared to significant increases with an IgG antibody control (*P* = .012) or no-antibody conditions (*P* = .0295) (Figure 1g) (Supplementary File 1A), indicating this integrin as the receptor for irisin on osteoclasts.

Next, we asked whether irisin-induced osteoclastogenesis led to enhanced bone resorption. Osteoclast differentiation cultures were performed on a variety of native and synthetic substrates with and without irisin (10 ng/mL). TRAP-positive osteoclasts were observed *in situ* on dentin slices after 7 days, and subsequent toluidine blue staining revealed a qualitative increase in resorption pit area on the surface with irisin treatment (Figure 2a). Irisin significantly increased osteoclast numbers on dentin (*P* = .013), as well as total resorption area (*P* = .045). When normalized by osteoclast number however, resorption was not significantly different (*P* > .99), indicating a dominant effect of cell number in increasing total resorption (Figure 2b). Irisin enhancement of total resorption was further confirmed via the Corning OsteoAssay, with a similar significant increase in total resorption area (*P* = .048) (Figure 2c). Release of carboxy-terminal collagen crosslinks (CTX) from osteoclast cultures on collagen substrates was measured to assess the effect of transient irisin treatment on resorption at earlier stages in culture, and was significantly increased with both continuous (*P* = .0153) and 24 hr (*P* < .0001) irisin exposure, with transient treatment also significantly higher than continuous (*P* = .0041) (Figure 2d) (Supplementary File 1B).

**Figure 2.**
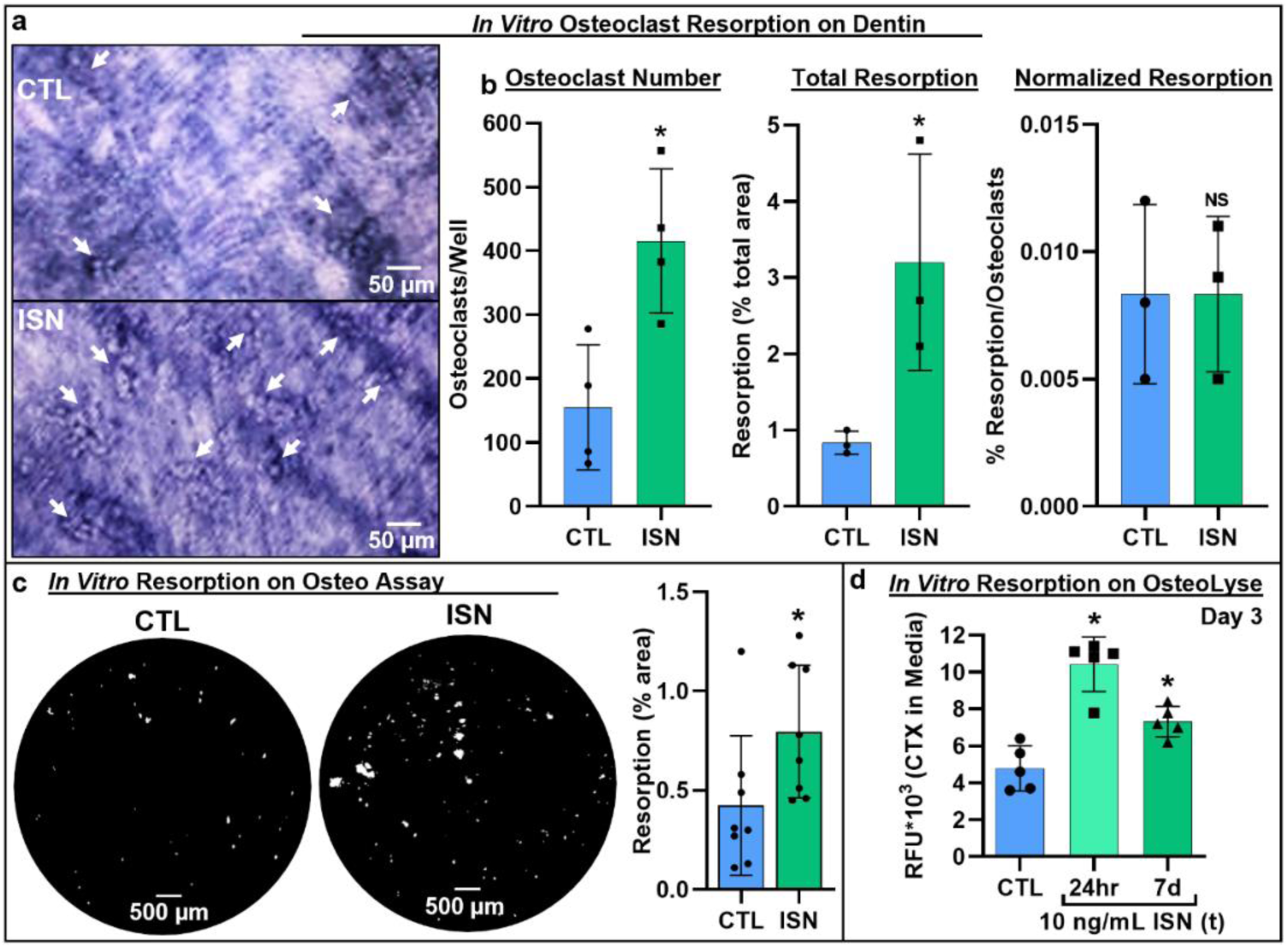
Representative images of resorption pits on dentin after 7-day osteoclast culture stained via toluidine blue dye, demonstrating both increased resorption with irisin treatment (ISN) versus untreated controls (CTL) **(a)**, with quantification of osteoclast number, total resorption area, and resorption normalized to osteoclast number **(b)**. Confirmation of irisin stimulation of resorption on Corning OsteoAssay calcium phosphate substrate, representative full-diameter images of 96 well plates with binary threshold to visualized resorption pits on Von Kossa-stained substrate after 7-day osteoclast culture for ISN vs. CTL, with quantification of total resorption by percent area **(c)**. Irisin stimulation early-stage resorption with transient treatment for first 24 hr alone, versus continuous ISN and CTL; quantification of resorption as determined by ELISA of CTX release into media, collected at day 3 of culture **(d)**. N = 4-8/group, \**P* < .05 vs. CTL.

To determine the key signaling pathways of irisin induced osteoclastogenesis we performed an unbiased analysis of RNA sequencing (RNAseq) data from irisin-treated and control hematopoietic progenitors, which demonstrated a qualitatively differential gene expression pattern as typified by hierarchical clustering, volcano plot, and principal component analysis (Figure 3a). Irisin treatment significantly increased expression of the resorption markers *Adamts5* (*P* = .0381) and *Loxl2* (*P* = .009), and markers for secreted clastokines known to stimulate osteoblasts: *Postn* (*P* = .0002), *Igfbp5* (*P* = .03), *Tgfb2* (*P* = .0073), and *Sparc* (*P* = .0365). Significant decreases in expression of the macrophage markers *Mst1r* (*P* = .0416) and *Itgax* (*P* < .0001) and the lymphocyte markers *Cd72* (*P* = .0086), *Slamf8* (*P* = .0052), and *H2-aa* (*P* = .0017) indicated a preferential shift of the hematopoietic progenitor lineage toward osteoclast differentiation (Supplementary File 1C). RT-qPCR further showed that irisin significantly increased expression of key differentiation markers, namely *Rank*, the receptor for RANKL (*P* = .019), *c-src* (*P* = .008), *Fam102a* (*P* = .039), *Nrf2* (*P* = .0008), and the osteoclast fusion markers *Dcstamp* (*P* = .0039) and *Atp6vod2* (*P* = .012). Other early markers of osteoclast differentiation such as *Cfos* (*P* = .41), *Itgb3* (*P* = .09), *Nfatc* (*P* = .11), *Rela* (*P* = .12), and *Rgs12* (*P* = .09) were slightly but not significantly increased. Significant increases were confirmed for both key resorption markers: *Acp5* (*P* = .012) and *Loxl2* (*P* = .035), and clastokine markers: *Sparc* (*P* = .009) and *Wnt10a* (*P* = .047) (Figure 3b) (Supplementary File 1D).

**Figure 3.**
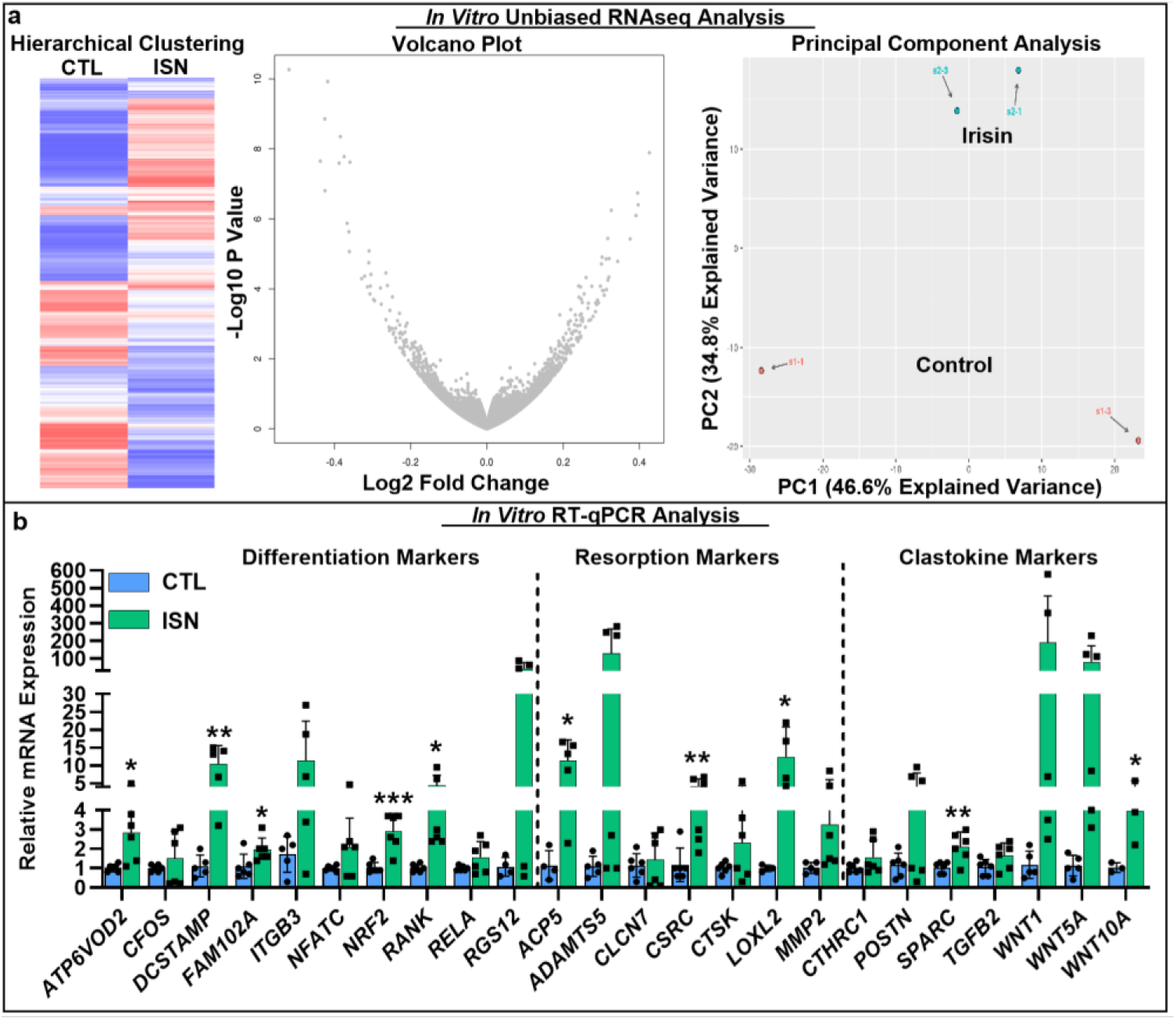
RNAseq analysis of differential gene expression patten induced irisin treatment (ISN) compared to untreated controls (CTL), as typified by representative sample hierarchical clustering, volcano plot, and principal component analysis **(a)**. Relative mRNA expression quantified by RT-qPCR of markers for osteoclast differentiation, resorption, and clastokines in irisin treated osteoclasts (ISN) compared to untreated controls (CTL), normalized to *Hprt* expression **(b)**. N = 3-6 samples/group. *P < 0.05, **P < 0.01, ***P < 0.001 vs. CTL within gene.

We next turned to a genetic model of forced expression of *Fndc5* in C57BL/6J mice using the muscle specific promoter *Mck*. At 4.5 months there was a marked reduction in cortical area, and a significant decrease in trabecular bone volume fraction compared to littermate controls (*P* = .012) (Figure 4a). Dynamic histomorphometry demonstrated similar decreases in overall bone volume fraction and trabecular thickness, and increased osteoclast numbers on the trabecular bone surface (Figure 4b). RT-qPCR analysis of whole bones demonstrated a significant increase in *Fndc5* (*P* < .0001), indicating promoter activity in the marrow in addition to muscle, and the osteoclast differentiation markers *Nfkb* (*P* = .0448) in transgenic versus wild type mice. Primary bone marrow progenitors from these mice had greater osteoclastogenic potential compared to wild type and yielded significantly higher numbers of osteoclasts that were qualitatively larger than controls during *in vitro* differentiation (*P* < .0001) (Figure 4d) (Supplementary File 1E).

**Figure 4:**
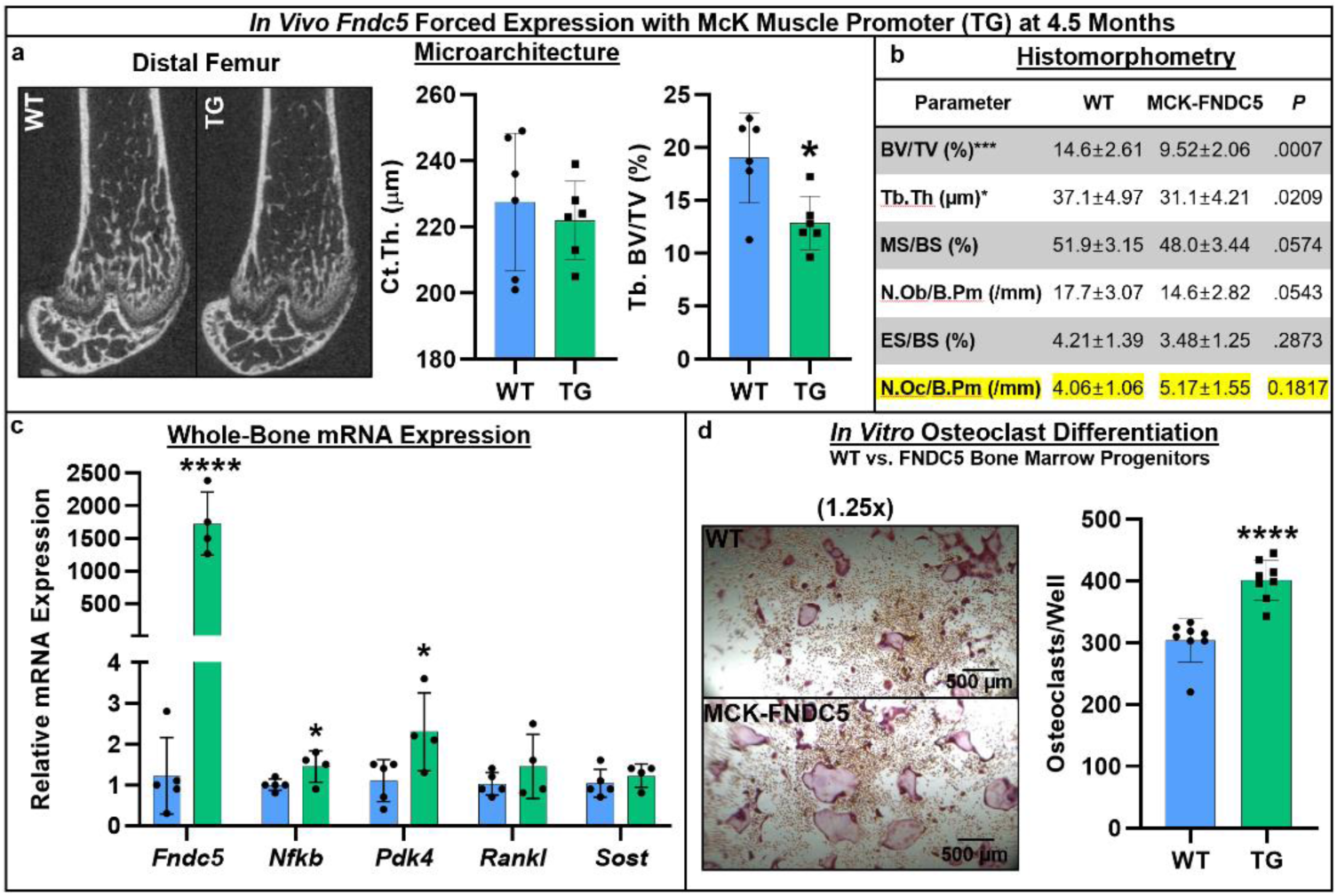
Skeletal phenotype and osteoclastogenic potential of *Fndc5* forced expression with McK muscle promoter mice (MCK-FNDC5) compared to wild type C57BL/6J controls (WT; N = 5/genotype). Representative distal femur microarchitecture at 4.5 months demonstrates qualitatively reduced cortical thickness, with quantification of significantly reduced trabecular and cortical bone parameters **(a)**. Tibial and vertebral histomorphometry at 4.5 months demonstrates continued reduction of bone volume fraction, with a non-significant increase in osteoclast numbers in the tibia **(b)**. While whole bone gene expression shows increased *Fndc5*, as well as promoters of osteoclast differentiation, *Nfkb, Pdk4, Rankl*, and *Sost* as normalized to *Hprt* **(c)**. In vitro MCSF/RANKL-induced osteoclast differentiation from bone marrow progenitors yielded higher osteoclast numbers in MCK-FNDC5 vs. WT **(d)**. N = 4-8, \**P* < .05, \*\*\**P* < .001, \*\*\*\**P* < .0001 vs. WT.

The present study demonstrates that irisin plays an important role in regulating bone remodeling not only by stimulating osteoblasts and osteocytes, but also by directly acting on osteoclasts to promote differentiation and resorption. This stimulatory effect was observed across multiple experiments with primary murine progenitors and the RAW 264.7 macrophage cell line and occurred with either intermittent or continuous irisin exposure across a range of physiologic concentrations previously reported in humans^1^. Analogous to its action on osteocytes, we confirmed expression of integrin subunits α_V_ and β_5_ expression on osteoclasts and identified it as a candidate receptor for irisin, particularly since blocking this receptor complex with a neutralizing antibody completely suppressed the stimulatory effect of irisin on osteoclastogenesis (Figure 1). Furthermore, we found that both recruitment and differentiation of more osteoclast progenitors with irisin treatment appeared to be the driving factors in enhanced bone resorption, based on *in situ* studies on native dentin as well as synthetic calcium phosphate and collagen substrates (Figure 2). Using unbiased RNAseq analysis and qRT-PCR of irisin-treated osteoclasts we noted that some markers of early osteoclast differentiation, nuclear fusion markers, and enzymes related to bone resorption were upregulated, matching the functional effects observed *in vitro*. In addition, several osteoclast-secretory factors known to stimulate osteoblasts were significantly upregulated, suggesting that the direct actions of irisin on this cell type may have further impact on cell signaling to enhance coupled remodeling (Figure 3).

To confirm the capacity of irisin to impact osteoclast function *in vivo*, we employed a genetic strategy with chronically forced expression of *Fndc5* using the muscle specific *Mck* promoter. Using that mouse model both *in vivo* and *in vitro* we demonstrated that high levels of irisin can drive osteoclastogenesis, although we cannot not exclude an indirect effect from osteocytic activation to drive resorption, despite the absence of increased *Rankl* or *Sost* expression (Figure 4). While further studies are necessary to fully elucidate the mechanisms of irisin actions on bone, the present work adds to previous studies demonstrating this myokine’s ability to both act directly on bone cells of distinct origin, and to modulate signaling from one cell type to one another. This may have major physiologic relevance since one acute effect of intense physical activity is a decrease in serum calcium, followed by a secondary rise in parathyroid hormone^9^. In the context of our experimental paradigm, it is conceivable that irisin represents another but more acute counter-regulatory hormone that works during the first minutes of exercise to tightly maintain serum calcium levels by its direct actions on osteoclasts and through osteocytic osteolysis. Taken together, our studies provide more evidence that irisin mediates muscle-bone cross talk by regulating bone remodeling.

## Methods

### Primary Osteoclast Culture

Primary murine osteoclasts were differentiated and cultured *in vitro* by the following methods. Bone marrow was collected via centrifugation from the femur and tibia of 8-week-old male C57BL6/J mice and cultured in αMEM (VWR, Radnor, PA, USA) supplemented with 10% fetal bovine serum VWR) and 1% Pen-Strep (VWR). After 48 hr, non-adherent hematopoietic progenitor cells were removed and plated at 1.563×10^5^ cell/cm^2^ in 96-well tissue culture plates for cell counting or in 12-well tissue culture plates for RNA extraction. Osteoclast differentiation was stimulated by supplementation with 30 ng/mL M-CSF (PeproTech, Rocky Hill, NJ, USA) and 100 ng/mL RANKL (PeproTech)^10^, with 200 µL or 2 mL media refreshed at days 3 and 5 after plating for 96 and 12 well plates respectively. Irisin was produced in HEK 293 cells as a 10 his-tag recombinant, via previously established protocols (Lake Pharma, Hopkinton, MA, USA)^6^, and supplemented continuously in the media at 10 ng/mL, or as otherwise indicated.

### Primary Source Gender and Cell Line Confirmations

8-week-old male C57BL6/J mice were used as the primary cell source for all experiments unless otherwise stated. Confirmation of irisin effect on osteoclastogenesis was also established in female C57BL6/J mice to compare gender among this wild type murine primary source of progenitors, which were cultured and counted as described. Additionally, the RAW 267.4 macrophage cell line (ATTC, Manassas, VA, USA) was employed as a non-primary cell source, following previously published protocols for osteoclast differentiation from this cell line^11,12^. Briefly, RAW cells were played at a lower density in 96-well plates of 6×10^3^ cells/well and cultured as described, but for the exclusion of MCSF in the media.

### Osteoclast Counts

At day 7, 96-well plates were fixed in 10% formalin and stained for TRAP (Acid Phosphatase Kit, Sigma-Aldrich, St. Lois, MO, USA) to visualize and count mature osteoclast numbers, where a TRAP-positive cell with 3 or more nuclei was defined as an osteoclast. Initial counting was performed via manual counting on an inverted microscope, with confirmative counts performed by manually counting blinded copies of composite images of each sample in ImageJ (Blind Analysis, Labcode).

### Integrin Antibody Blocking

A separate experiment employed the culture and counting methods described above for irisin treatment in combination with a neutralizing antibody for integrin α_V_β_5_ (Anti-Integrin aVb5, MilliporeSigma, Burlington, MA, USA) and an IgG control (Mouse IgG1 Isotype Control, R&D Systems, Minneapolis, MN, USA), each supplied continuously in the media during the 7 day culture at 0.9 µg/ml.

### Resorption Assays

Multiple resorption assays were utilized to characterize and confirm the effect of irisin treatment on osteoclast resorptive capacity. The primary approach employed decellularized dentin slices as a native bone substrate. Hematopoietic progenitors were plated on the bone slices at 3×10^5^ cell/cm^2^ in a 50 µL place on top of slice in a 10 mm petri dish and incubated for 30 minutes to facilitate cell adhesion to the substrate alone. Slices were then moved into 96-well plates and cultured as described above. At day 7, dentin slices were TRAP stained and imaged as described for osteoclast counts, then sonicated briefly to removed cells and stained with toluidine blue to visualize resorption pits by previously published methods^13^. Briefly, each slice was placed face-down on a 20 µL drop of 1% toluidine blue solution (in 1% sodium borate 10-hydrate solution with distilled water) for four minutes and then rinsed and allowed to airdry prior to imaging and manual calculation of total pit areas for blinded images in ImageJ. Confirmation experiments were performed with the OsteoAssay resorption assay (Corning Inc., Corning, NY, USA), whereby osteoclasts were cultured on the substrate treated 96-well plates by the previously described methods, then removed with 10% bleach and the substrate was stained with Von Kossa stain (Sigma-Aldrich). Imaging of the wells on a dissecting scope with backlighting allowed visualization of pits, and automated area calculations based on binary thresholded masks of the images. To determine earlier time-point resorption, the OsteoLyse assay (Lonza, Basel, Switzerland) was employed, utilizing the same osteoclast culture methods on a collagen substrate, whereby detection of carboxy-terminal collagen crosslinks (CTX) allows for relative quantification of degraded collagen as an indicator of resorptive activity. Aliquots of the media at day 3 in culture were analyzed via ELISA for relative fluorescence indicative of CTX release and normalized to undifferentiated and no-cell controls.

### Gene Expression Analysis

Total RNA was isolated from osteoclast cells at day 7 with the Trizol reagent (ThermoFisher, Waltham, MA) and RNeasy mini kit (Qiagen, Hilden, Germany), with mRNA enrichment from 100 ng of total purified RNA and Illumina sequencing libraries preparation performed using Kapa stranded mRNA Hyper Prep (Roche Sequencing Solutions, Pleasanton, CA, USA). Gene libraries were multiplexed in an equimolar pool and were sequenced on an Illumina NextSeq 500 with single-end 75 bp reads. Raw reads were aligned to the UCSC mm10 reference genome using a STAR aligner^14^ (version STAR_2.4.2a), and raw gene counts were quantified using the quantMode GeneCounts flag. Differential expression testing was performed using Limma^15^ and DESeq2^16^. RNAseq analysis was performed using the VIPER snakemake pipeline^17^. Follow-up RT-qPCR was performed on RNA from a separate set of osteoclasts for markers identified via RNAseq and additional targets for differentiation, resorption, and clastokines with primer sequences obtained PrimerBLAST (NCBI-NIH), using reverse transcriptase kit (Qiagen) and AzuraQuant Green Fast PCR Mix (Azura Genomics, Rynam, MA, USA) with an IQ PCR detection system (Bio-Rad, Hercules, CA, USA). Gene expression data were analyzed via the comparative Ct method, utilizing *Hprt* as the housekeeping gene and normalizing by untreated control mean Ct.

### Forced Expression of Fndc5 in Murine Muscle

Transgenic C57BL/6J mice with forced expression of *Fndc5* were generated and generously gifted by Dr. Eric Elson of UT Southwestern. The *Mck* promoter was utilized as previously demonstrated to induce skeletal muscle-specific forced expression of *Fndc5*^*18*^. Microarchitecture of distal trabecular bone and midshaft cortical bone was analyzed at 4.5 months by µCT and static and dynamic histomorphometry, with measures performed and analyzed according to standard nomenclatures. Additional femur and tibia were pulverized, and RNA extraction and subsequent gene expression analysis was performed via aforementioned protocols. Bone marrow isolation and *in vitro* osteoclast cultures were performed via previously described protocols. All experiments were conducted with 6 age-matched female mice.

### Experimental Design and Data Analysis

Isolation of primary murine bone marrow was conducted by pooling tissue from the maximum available number of same-gender littermates (3-5). For *in vitro* cultures, the adequate number of biological replicates (replicate wells in a tissue culture plastic plate, or slices of dentin) was determined via power analyses based on preliminary data (α = .05, Power = 0.8) as 6, and so osteoclast counting and resorption experiments were conducted in triplicate with representative experiments shown, with 6-8 replicate wells per group in each experiment. Similarly, gene expression analysis via RT-qPCR was conducted for duplicate repeat experiments with three biological replicates per group and two technical replicates (replicate wells read per sample and averaged), with pooled representative data for a sample number of 6. Due to limitations in the availability of dentin slices, this resorption experiment was conducted once with a sample size of 5 per group. Similarly, RNA sequencing analysis was performed on a separate experimental set with three biological replicates per group. For characterization of the *Fndc5*-transgenic mouse, power analyses based on previous outcome metrics from histomorphometry and gene expression in the Fndc5-null mouse experiments^6^ (α=0.05, Power=0.8) indicated a n adequate sample size of 6 mice per group, which was employed for *in vivo* characterization of bone properties based on the availability of same-gender age-matched mice, while i*n vitro* culture of osteoclast progenitors from these mice were carried out with bone marrow isolates from maximal number of same-gender littermates (3-5) and 8 replicate wells of osteoclast differentiation cultures per group.

For statistical analysis, outlier identification was first performed via Grubb’s test with α = .05. Based on recommended guidelines for analysis, comparisons between two groups alone (irisin vs. control osteoclast counts, resorption, gene expression) was conducted via unpaired, two-tailed t-test, *P* < .05, while multiple group comparisons were made via ordinary one-way (irisin dose and duration versus control) or two-way (antibody/irisin treatment osteoclast counts) ANOVA with Tukey post-hoc analysis and *P* < .05. RNA sequencing data analysis was performed as described above with statistical significance via Wald test with Benjamini-Hochberg adjustment. Quantitative data are represented graphically as mean ± standard deviation with individual values overlaid.

## Acknowledgments

This work was funded in part by NIH/NIAMS F32AR077382, NIH U19AG060917, NIH U54GM115516-01A1, NIH/NIDDK R01 DK112374, and NIH/NIGMS 1P20GM121301. We thank Zach Herbert, Maura Berkeley and Andrew Caruso from the Molecular Biology Core Facilities at the Dana-Farber Cancer Institute for RNAseq. We thank Dr. Eric Elson of UT Southwestern for providing the transgenic *Fndc5* mice.

## Competing Interests

The authors have no conflicts of interests to disclose.

